# Learning is shaped by abrupt changes in neural engagement

**DOI:** 10.1101/2020.05.24.112714

**Authors:** Jay A. Hennig, Emily R. Oby, Matthew D. Golub, Lindsay A. Bahureksa, Patrick T. Sadtler, Kristin M. Quick, Stephen I. Ryu, Elizabeth C. Tyler-Kabara, Aaron P. Batista, Steven M. Chase, Byron M. Yu

**Author notes:** These authors contributed equally to this work (Aaron P. Batista, Steven M. Chase, Byron M. Yu).

## Abstract

Internal states such as arousal, attention, and motivation are known to modulate brain-wide neural activity, but how these processes interact with learning is not well understood. During learning, the brain must modify the neural activity it produces to improve behavioral performance. How do internal states affect the evolution of this learning process? Using a brain-computer interface (BCI) learning paradigm in non-human primates, we identified large fluctuations in neural population activity in motor cortex (M1) indicative of arousal-like internal state changes. These fluctuations drove population activity along dimensions we term neural engagement axes. Neural engagement increased abruptly at the start of learning, and then gradually retreated. In a BCI, the causal relationship between neural activity and behavior is known. This allowed us to understand how these changes impacted behavioral performance for different task goals. We found that neural engagement interacted with learning, helping to explain why animals learned some task goals more quickly than others.

## Introduction

As we move about the world, we experience fluctuations in internal states such as arousal, motivation, and engagement. These states are governed by the modulation of neural activity throughout the brain (Aston-Jones and Cohen, 2005; McGinley et al., 2015; Allen et al., 2019; Stringer et al., 2019; Steinmetz et al., 2019). The manner in which these modulations relate to the ongoing computations performed by the cerebral cortex is not well understood. In predominantly sensory areas of cortex, changes in an animal’s internal state are known to affect neural response magnitude, signal-to-noise ratio, timing, and variability (Luck et al., 1997; Mitchell et al., 2007; Cohen and Maunsell, 2009; Noudoost et al., 2010; McGinley et al., 2015; Vinck et al., 2015). Depending on how these changes align with respect to neural encoding of stimulus information or downstream readout, changes in an animal’s internal state can impact perceptual processing and decision making (Averbeck et al., 2006; Moreno-Bote et al., 2014; Ruff and Cohen, 2019; Cowley et al., 2020). Changes in internal state are also known to impact motor control and behavior, as the speed and latency of both eye movements and arm reaches are known to be modulated by signals such as motivation, intrinsic value, and reward expectation (Sugrue et al., 2004; Mazzoni et al., 2007; Xu-Wilson et al., 2009; Leathers and Olson, 2012; Dudman and Krakauer, 2016; Sedaghat-Nejad et al., 2019; Shadmehr et al., 2019). These studies and others illustrate the importance of understanding the influence of internal states on sensory processing and behavior.

What has been less well studied is the impact of internal state changes on learning (Figure 1A). When we learn to perform a task, such as shooting a basketball, the firing activity of populations of neurons in the brain (Figure 1A, gray clouds) is modified in a particular manner in order to drive improved behavior (Figure 1A, blue and red clouds) (e.g., Li et al. (2001); Andalman and Fee (2009); Keller and Hahnloser (2009); Ganguly and Carmena (2009); Gu et al. (2011); Koralek et al. (2012); Hwang et al. (2013); Jeanne et al. (2013); Law et al. (2014); Sadtler et al. (2014); Poort et al. (2015); Athalye et al. (2018); Golub et al. (2018); Vyas et al. (2018); Perich et al. (2018); Oby et al. (2019)). We also know that while animals perform a task, neural activity undergoes internal state fluctuations that are not directly related to task performance (Figure 1A, orange arrows) (e.g., Arieli et al. (1996); Cohen and Maunsell (2009); Churchland et al. (2010); Ecker et al. (2014); Schölvinck et al. (2015); Lin et al. (2015); Rabinowitz et al. (2015); Ni et al. (2018); Stringer et al. (2019); Cowley et al. (2020)). Depending on the task goals, changes in internal state have the potential to make some learning-related neural changes easier to achieve (Figure 1A, blue cloud), while other changes may be made more difficult (Figure 1A, red cloud). When changes due to internal state are incongruous with learning, how do neural populations modify their activity to drive improved behavior? One possibility is that the internal state fluctuations that make learning more difficult might be suppressed. Alternatively, the impact of internal state fluctuations on learning may be unavoidable, which may result in some task goals being harder to achieve than others.

**Figure 1.**
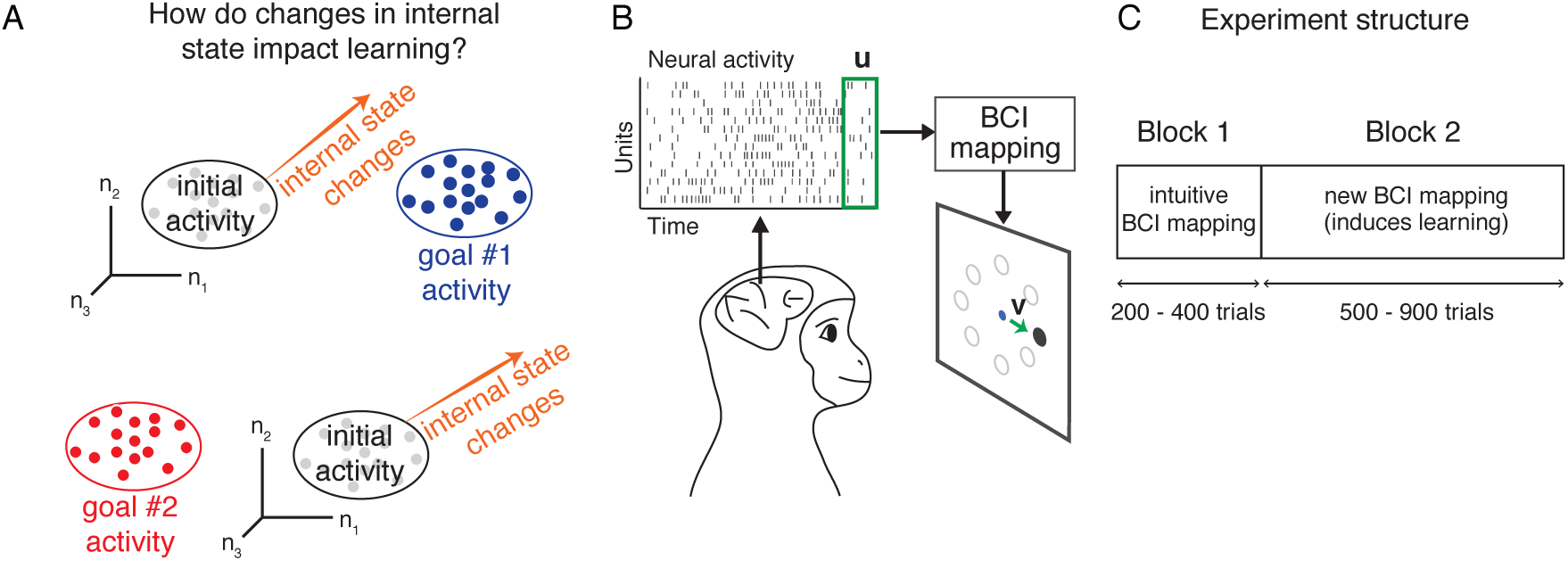
Studying how changes in neural activity during learning are impacted by changes in internal state. **A.** Here we ask whether changes in internal state impact how neural population activity is modified during learning. Before learning, neural activity resides in some region (‘initial activity’) of population activity space, depicted here by the firing rates of three neurons (*n*_1_, *n*_2_, *n*_3_). During learning, the neural activity needs to migrate to a different region of population activity space to achieve a particular task goal (‘goal #1 activity’ and ‘goal #2 activity’). Changes in the animal’s internal state can push the neural activity closer to (top orange arrow) or further from (bottom orange arrow) the region appropriate for achieving a given task goal. **B.** Monkeys performed an eight-target center-out task using a brain-computer interface (BCI). Neural activity was recorded using a multi-electrode array implanted in M1. Spike counts (**u**) were taken in 45 ms bins (green box). The BCI mapping converted the neural activity into a cursor velocity (**v**) at each 45 ms time step, updating the position of a visual cursor on a screen. Monkeys were rewarded for successfully guiding the cursor to hit the visually instructed target. **C.** Each experiment consisted of two blocks of trials. In Block 1, a monkey completed 200-400 trials using an intuitive BCI mapping. In Block 2, the monkey completed 500-900 trials with a new BCI mapping he had not used before.

Answering this question is challenging because the causal relationship between neural activity and behavior is not known in general. This makes it difficult to understand which changes to neural activity would yield improved performance, as well as how fluctuations in internal state would either interfere or augment that performance. To address this difficulty we can leverage a brain-computer interface (BCI) (Taylor et al., 2002; Carmena et al., 2003; Hochberg et al., 2006; Ganguly and Carmena, 2009; Gilja et al., 2012; Hauschild et al., 2012; Sadtler et al., 2014), where the causal relationship, or ‘mapping,’ between neural activity and behavior is known exactly and determined by the experimenter.

We trained three rhesus monkeys to modulate the activity of ∼90 units in primary motor cortex (M1) to move a computer cursor on a screen using a BCI (Sadtler et al., 2014). In previous work, we compared the neural population activity before versus after monkeys learned to use a new BCI mapping (Golub et al., 2018; Hennig et al., 2018). Here we study how neural activity changed throughout learning, and the degree to which these changes were influenced by fluctuations in the monkey’s internal state.

We first identified the dimensions of the largest fluctuations in M1 population activity. Surprisingly, abrupt changes in population activity along these dimensions were triggered by changes in various aspects of the task, ranging from brief pauses in the task to perturbations of the BCI mapping. Furthermore, trial-to-trial changes in population activity along these dimensions were correlated with changes in the monkey’s pupil size. These observations suggested that changes in population activity along these dimensions could be related to the monkey’s arousal, engagement with the task, or motivation throughout the experiment. For this reason, we term these dimensions *neural engagement* axes.

To induce learning, we perturbed the mapping between neural activity and cursor movements, requiring monkeys to modify the neural activity they produced in order to restore proficient control of the cursor towards each target. This allowed us to study how changes in activity along neural engagement axes interacted with learning. We found that neural population activity did not take a direct path from the activity produced prior to learning to the activity produced at the end of learning. In particular, neural activity changed abruptly along the neural engagement axes at the start of learning. This change occurred regardless of the relationship between neural engagement axes and cursor movements, which led to an immediate improvement in performance for some targets and impaired performance for others. Following the abrupt change, neural activity retreated along neural engagement axes, which interacted with learning. This led to monkeys learning some targets more quickly than others, in a predictable manner based on how neural engagement interacted with the demands of the learning task. These results indicate that changes in internal states can influence how quickly different task goals are learned.

## Results

To understand how changes in internal state might interact with learning (Figure 1A), we trained three monkeys to perform an eight-target center-out task using a brain-computer interface (BCI) (Figure 1B; see Methods). On each trial, monkeys controlled a computer cursor by modulating neural activity recorded from primary motor cortex (M1). The relationship between the recorded neural activity and cursor velocity was specified by the BCI mapping. In each experimental session, monkeys used two different BCI mappings (Figure 1C). During the first block of trials, monkeys used an ‘intuitive’ BCI mapping, calibrated so as to provide the monkey with proficient control of the cursor. After monkeys performed the task for a few hundred trials using the intuitive mapping, we changed the mapping between neural activity and cursor movement to a new BCI mapping that the monkey had not used before (a ‘within-manifold perturbation’; see Sadtler et al. (2014)).

Prior to each experiment, we applied factor analysis (FA) to identify the top ten dimensions, or factors, capturing the most covariability of the neural population activity. The BCI mappings presented during each experiment were chosen such that the cursor velocity was determined by only these top ten factors. In order to ensure that our results captured changes in neural activity describing substantial covariance in the population, here we analyze neural activity only in these top ten factors.

### Neural population activity in primary motor cortex modulates with monkeys’ engagement

We first show that the neural population activity recorded during these experiments reflected a correlate of the monkey’s internal state. We observed that, while monkeys used the intuitive mapping, the neural activity produced for a given target showed substantial trial-to-trial variability (Figure 2A, gray dots). We found the direction of greatest variance of the neural activity for each target (Figure 2A, orange line). Surprisingly, later in the session when the new BCI mapping was introduced, neural activity on the first trial to a given target showed an abrupt change from the average neural activity during Block 1, with this change occurring almost directly along the axis identified earlier (Figure 2B, compare ‘1st trial of Block 2’ to ‘avg. during Block 1’). Interestingly, on subsequent trials, neural activity gradually retreated down this same axis (Figure 2B, grayscale indicates trial index).

**Figure 2.**
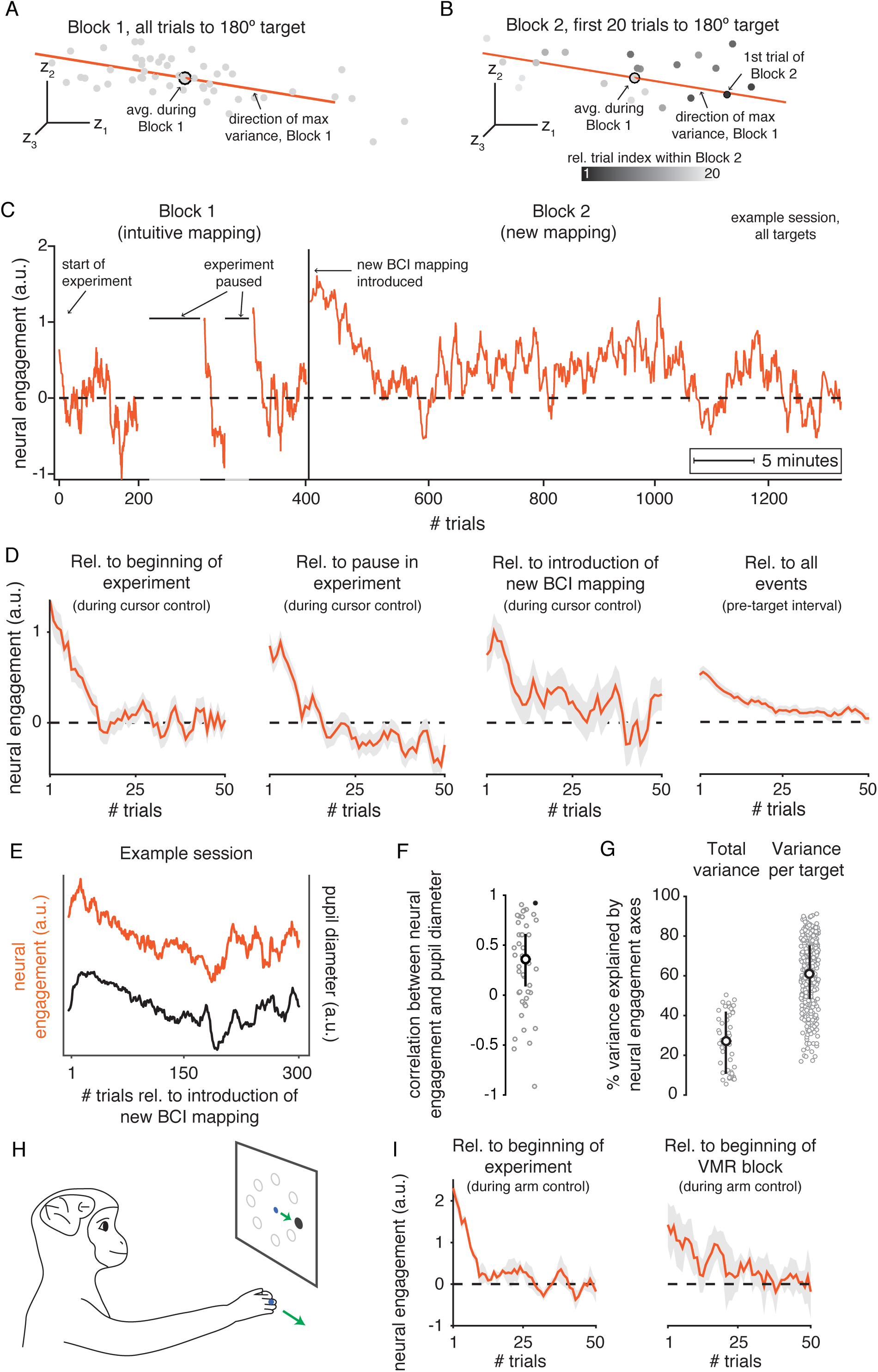
Neural activity increased abruptly along a neural engagement axis following experimental events. **A.** Neural activity in the top three factor dimensions of highest covariance (*z*_1_, *z*_2_, *z*_3_) for trials to the same target from Block 1 of session J20120528. Each gray point is the average neural activity during a single trial. Orange axis depicts the direction of maximum variance of all gray points. The axis was defined in the 10-dimensional factor space, although only the top three dimensions are depicted here. **B.** Same as A, but for the first 20 trials to the same target during Block 2 (color indicates trial index). Orange axis from A shown for reference. *Neural engagement* for each trial is the projection of neural activity onto the axis identified during Block 1 for trials to the same target. **C.** Neural engagement over time from session J20120528, with annotations indicating timing of various events controlled by the experimenter. Position along horizontal axis indicates clock time (see legend), with trial indices marked for reference. Horizontal dashed line at zero indicates average neural engagement during Block 1 (see Methods). **D.** First three subpanels: Neural engagement averaged across sessions from all monkeys during cursor control relative to the start of the experiment, the longest pause during Block 1, and the start of Block 2. Last subpanel: Neural engagement during the interval of each trial before the monkey had seen the target (see Methods), averaged across all three experimental events. Shading indicates mean ± SE across sessions. **E.** Neural engagement during Block 2 from example session shown in C, alongside monkey’s average pupil size during the same trials. **F.** Pearson’s correlation between neural engagement and pupil size during Block 2 for all sessions (dots), with session from E indicated in black. White circle and black lines depict the bootstrapped median and 95% C.I. of the correlations across sessions, respectively. **G.** Percentage of shared covariance of neural population activity explained by neural engagement axes, when including trials to all targets in a session (‘Total variance’), or only trials to a single target (‘Variance per target’). White circle depicts median; error bar depicts median ± 25^*th*^ percentile of correlations across sessions. **H.** In a different set of experiments, a monkey performed a center-out task by moving its hand to control the cursor’s position (see Methods). **I.** Neural engagement averaged across sessions from hand control experiments, both relative to the beginning of the experiment (left), and relative to the introduction of a visuomotor rotation (right). Same conventions as D.

We next quantified how these trial-to-trial changes in neural activity progressed throughout the experiment (Figure 2C). To do this, we identified the axis of greatest variability during Block 1 for each target separately (e.g., the orange axis in Figure 2A-B), and projected the neural activity for each trial along the appropriate target-specific axis. So that we could compare these values across trials to different targets, we z-scored the projected values for each target separately (see Methods). This yielded a trial-by-trial measure we will refer to here as *neural engagement*, for reasons we discuss below.

Neural engagement abruptly increased and gradually decreased following various experimental events, beyond just the introduction of the new BCI mapping (Figure 2C). For example, neural engagement was initially elevated on the very first trials of the experiment, and then gradually decreased on later trials (Figure 2C, “start of experiment”). Next, near the middle of Block 1, the experimenter would pause the experiment for a few minutes to choose the BCI mapping that would be introduced in the upcoming Block 2. Following these pauses (Figure 2C, “experiment paused”), neural engagement increased, and then gradually subsided. Finally, a few minutes later when the experimenter seamlessly introduced the new BCI mapping (without pausing the experiment), neural engagement again abruptly increased (Figure 2C, “new BCI mapping introduced”) and gradually subsided on subsequent trials. We observed similar neural changes across multiple sessions from all three monkeys (Figure S1), indicating that these changes were not specific to the particular BCI mappings used during a given session. Rather, these changes in neural activity appeared to reflect generalized changes in the monkey’s internal state throughout the experiment, and could reflect changes in arousal (Vinck et al., 2015), engagement with the task (Steinmetz et al., 2019), or motivation (Mazzoni et al., 2007). While the specific source of these changes is as yet unknown (we discuss various possibilities in Discussion), these changes have important consequences for learning.

Two additional aspects of neural engagement are consistent with it reflecting variations in the monkey’s internal state. First, when averaged across all sessions, neural engagement showed a consistent time course following each experimental event: an immediate increase on a single trial, followed by gradual decay over subsequent trials (Figure 2D). These changes in neural engagement appeared not only during the period within each trial while the monkey was controlling the cursor (Figure 2D, first three panels), but also during the beginning of each trial before the monkey had seen the visual target (Figure 2D, last panel). Thus, neural engagement remained elevated even when the monkey was not actively performing the task, consistent with this signal reflecting a slowly-varying change in the monkey’s internal state. Second, changes in an organism’s internal state are typically correlated with changes in its pupil size (McGinley et al., 2015). In agreement with this, we found that fluctuations of neural engagement were often strikingly positively correlated with changes in the monkey’s pupil size (Figure 2E). Across sessions, the median Pearson’s correlation between neural engagement and pupil size was *ρ* = 0.36 (bootstrapped 95% C.I. [0.10, 0.60]) (Figure 2F), similar to levels observed in other work (Cowley et al., 2020).

Changes in activity along the neural engagement axes accounted for a substantial amount of the covariance of the population activity. When considering population activity during Block 1 across trials to all eight targets—and thus also including the across-target variance in neural activity due to the monkey aiming towards different targets—changes in neural engagement explained ∼30% of the total trial-to-trial variance of the factor activity (Figure 2G, “Total variance”). Within trials to the same target, changes along the neural engagement axis explained ∼60% of the trial-to-trial variance (Figure 2G, “Variance per target”). These results indicate that the trial-to-trial changes in population activity along the neural engagement axes were substantial.

To assess whether similar changes in neural engagement were present during arm movements (as opposed to BCI control), we analyzed data from a fourth monkey performing an eight-target center-out task by controlling a computer cursor with his hand (Figure 2H; see Methods). As with the BCI experiments, we identified a set of neural engagement axes in the population activity after applying factor analysis. We found that neural engagement was elevated both at the beginning of each experiment, and following the introduction of a visuomotor rotation (Figure 2I), with a time course that was strikingly similar to that of BCI control (Figure 2D). Taken together, we found that neural population activity in M1 during both BCI control and hand control showed large, trial-to-trial variations with a consistent time course relative to experimental events. In the following, we focus on BCI control, where we know the causal relationship between neural activity and behavior. This enables us to directly assess how changes in neural engagement relate to behavior (i.e., cursor movements).

### Studying the impact of changes in neural engagement on behavior using a BCI paradigm

Having established the presence of large fluctuations in neural engagement in M1, we next wanted to understand how these fluctuations might impact learning. Specifically, we sought to understand how the monkey’s ability to learn to move the cursor in a given direction with the new BCI mapping might be impacted by the relationship between the neural engagement axes and the new mapping.

First, we explain how a BCI paradigm allows us to quantify the interaction between neural engagement and behavior (i.e., cursor velocities). Consider a schematic of the neural activity produced by the monkey during Block 1 (Figure 3A, left subpanel). For trials to a given target (e.g., the 180° target), we can summarize the monkey’s average neural activity as a point in neural space (**z**, gray sphere), where here we depict the neural activity in the three factor dimensions of highest variance. The average cursor velocity under the intuitive BCI mapping (**v**, gray circle, top right subpanel) is given by projecting the neural activity onto the intuitive BCI mapping (**v** = *M*_1_**z**). During Block 1, the monkey’s average cursor velocities were near the target direction (Figure 3A, gray dashed line in top right subpanel), indicating the monkey’s ability to produce cursor movements that moved the cursor towards the target on average. We can also characterize the effect of an increase in neural engagement on cursor velocities by projecting the neural engagement axis (Figure 3A, orange arrow in left subpanel) onto the intuitive BCI mapping (Figure 3A, orange arrow in top right subpanel). In this case, increased neural engagement would result in a faster cursor speed towards the target.

**Figure 3.**
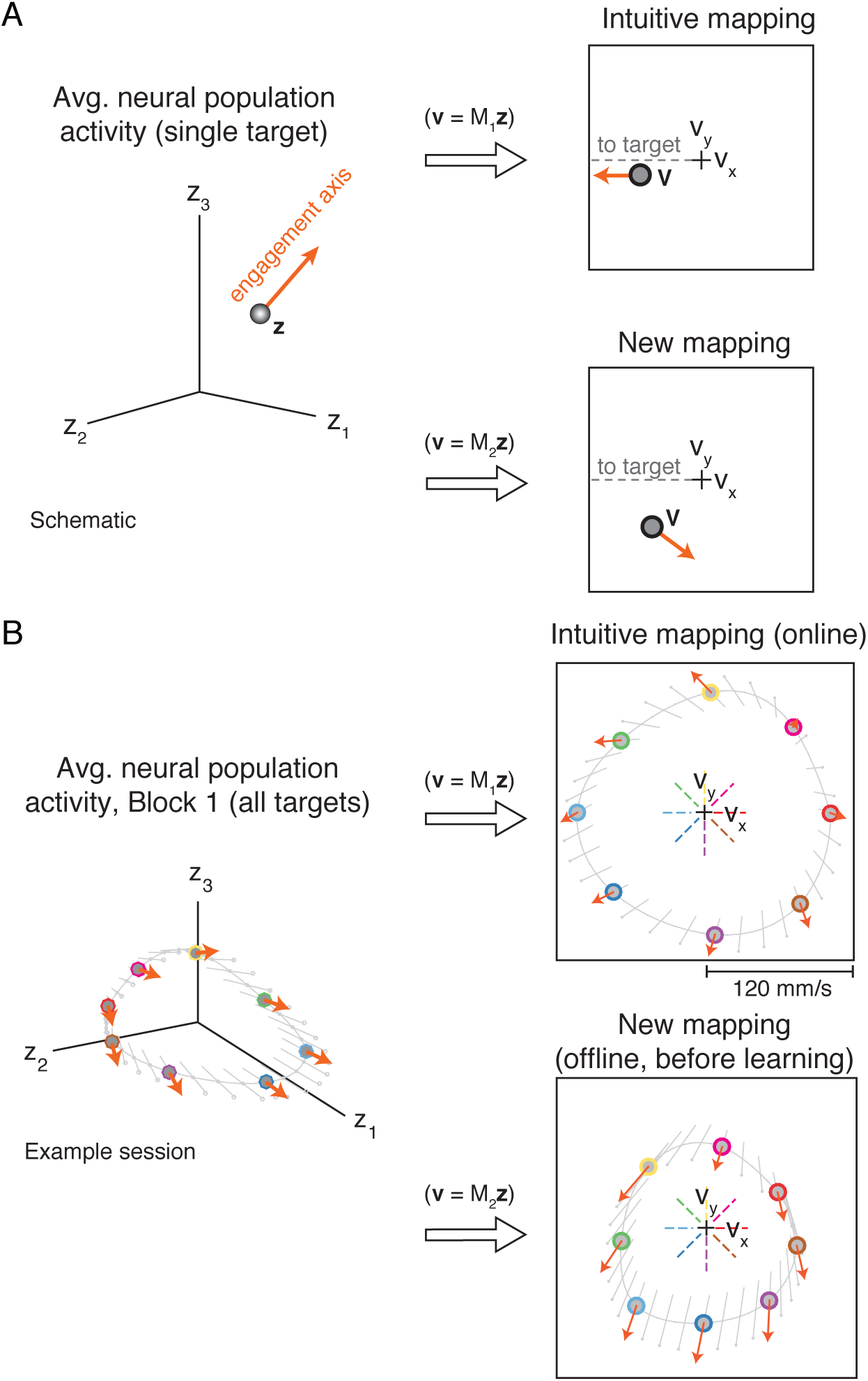
Predicting the impact of neural engagement on behavior during a BCI learning task. **A.** Left: Schematic of the average neural activity (**z**) recorded for trials to the same target during Block 1, along with the direction in which this activity is expected to move following an increase in neural engagement (orange arrow). Top right: Using the intuitive BCI mapping (*M*_1_), we can inspect the intuitive cursor velocity (**v**, gray circle) corresponding to **z**, as well as how this velocity will change if neural engagement increases (orange arrow). In this case, increased neural engagement will result in faster cursor movements towards the target (gray dotted line). Zero velocity is indicated by the black cross. Bottom right: We can repeat the same procedure using the new BCI mapping (*M*_2_) with the same neural activity **z** and neural engagement axis. **B.** For an example session, the average neural activity (gray circles with colored outlines) and engagement axes (orange arrows) for all eight targets. Gray lines indicate interpolations between the neural engagement axes for each target. Dashed colored lines in the two right subpanels indicate the eight target directions. Extent of box is ± 120 mm/s.

Next, consider the first trial of Block 2, when the monkey first encounters the new BCI mapping. If the monkey were to continue to produce the same average neural activity that he did during Block 1 (Figure 3A, left subpanel), this would no longer result in cursor movements straight to the target (Figure 3A, bottom right subpanel). Thus, the monkey must learn how to modify the average neural activity he produces in order to produce faster cursor speeds in the target direction. Importantly, the new BCI mapping also changes the manner in which neural engagement relates to cursor velocity. For this target, increasing neural engagement will move the cursor velocities even further from the target direction (Figure 3A, bottom right subpanel, orange arrow). In this manner, changes in neural engagement will interact with the monkey’s attempts to move the cursor towards the target.

We can gain a more holistic picture of the interaction between neural engagement and cursor velocities by visualizing the neural activity produced for all eight targets together (Figure 3B), shown here for an example session. We observed that, when visualized in factor space (Figure 3B, left subpanel), the neural engagement axes identified for different targets often appeared quite similar. In fact, across all targets and sessions, neural engagement axes were almost always consistent with the firing rates of all neural units changing in the same direction (Figure S2). Because of this, along with the manner in which we identified the sign of each neural engagement axis (see Methods), increases in neural engagement corresponded to increased firing rates in nearly all units. However, while the neural engagement axes for different targets were similar in terms of how they related to single unit firing rates, these axes also showed behaviorally relevant differences. For example, under the intuitive BCI mapping, increases in neural engagement sometimes led to faster speeds towards each target (Figure 3B, top right subpanel). A similar feature was also present during arm movements: After identifying the linear mapping of neural population activity most predictive of ensuing hand velocities, increases in neural engagement typically predicted faster hand speeds towards each target (Figure S3). These target-specific relationships between neural engagement axes and velocity indicate that changes in neural engagement impacted neural population activity differently depending on the direction in which the monkey was intending to move.

We now focus on the velocities under the new BCI mapping, as this indicates the initial cursor velocities the monkey would expect to produce during Block 2, were he to continue producing the same activity he did during Block 1. As discussed above, neural engagement can have different effects on cursor velocities depending on the direction in which the monkey is trying to move the cursor. In particular, increased neural engagement may lead to increased speeds towards some targets (e.g., Figure 3B, purple target in bottom right subpanel) and decreased speeds towards other targets (e.g., Figure 3B, pink target in bottom right subpanel). Additionally, increased neural engagement can affect not just the speed but also the direction of the velocity, leading to either decreased or increased angular error relative to the target direction (e.g., red and yellow targets, respectively). Overall, we observed that the new BCI mappings induced a variety of different relationships between neural engagement and cursor velocity, both across sessions and within targets of the same session (Figure S4). Thus, these experiments provided us with the means to assess how the different relationships between neural engagement and cursor velocity might impact the manner in which these different targets were learned.

### Neural engagement increased initially regardless of its impact on performance

To study the impact of changes in neural engagement on learning, we first characterized the level of neural engagement on the very first trial to each target using the new BCI mapping. As shown earlier, monkeys’ initial reaction to the introduction of the new mapping was, on average, to increase neural activity along the neural engagement axis (Figure 2D, third panel). However, as we have also shown, there are a variety of ways in which the neural engagement axes affected velocities under the second mapping (Figure 3C). This raises the possibility that neural engagement might have increased more for some targets than for others, depending on whether increasing neural engagement was expected to increase (Figure 4A) or decrease (Figure 4B) the speed of the cursor towards the target under the new mapping.

**Figure 4.**
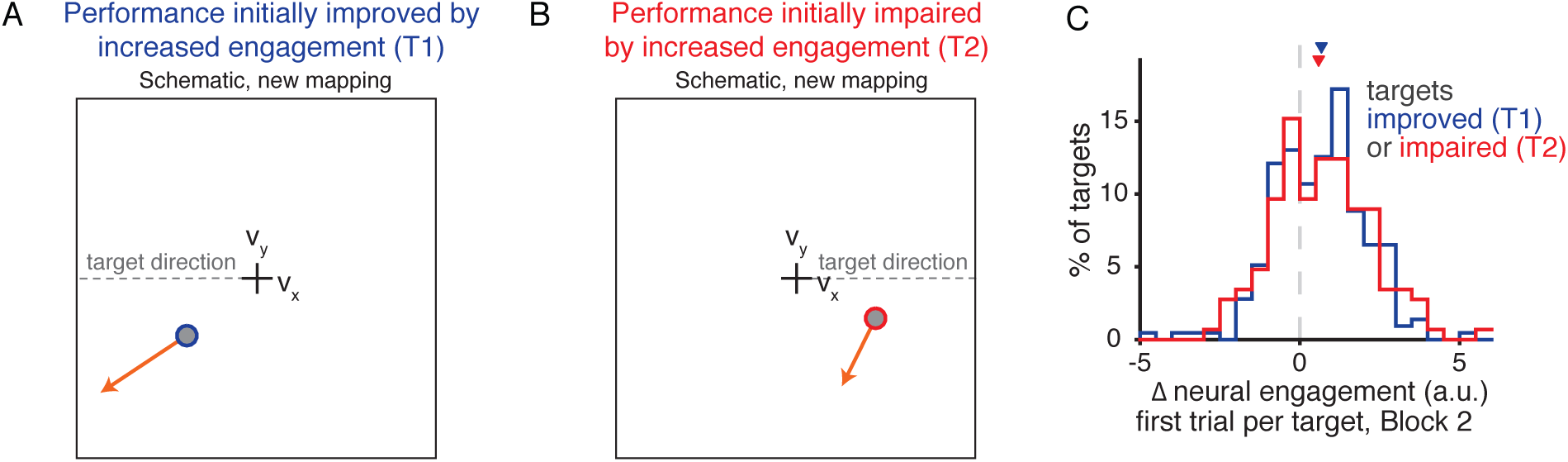
Neural engagement increased on the first trial of a learning task regardless of its impact on task performance. **A.**-**B.** Schematics depicting how increased neural engagement can lead to either faster (A) or slower (B) cursor speeds towards the target direction under the new BCI mapping. Same conventions as bottom right panel of Figure 3A. **C.** Distribution of the increase in neural engagement on the first trial to each target during Block 2, as a function of whether performance under the new mapping was expected to be improved (blue) or impaired (red) by an increase in neural engagement (as in A-B). Triangles depict the median of each distribution.

We anticipated that neural engagement would increase more for targets where doing so resulted in faster cursor speeds towards the target. To assess whether this was the case, we used the average activity from Block 1 to estimate the average expected velocity under the new mapping (Figure 4A-B, filled circles), as well as the expected impact on that velocity if neural engagement increased (Figure 4A-B, orange axes). We then classified each target as belonging to one of two groups, based on whether an increase in neural engagement was expected to increase (‘T1’, Figure 4A) or decrease (‘T2’, Figure 4B) the speed of the cursor towards the target direction.

We next assessed the levels of neural engagement on the first trial to each target in Block 2. Surprisingly, across targets from all sessions, the distribution of neural engagement on the first trial using the new mapping did not differ as a function of how performance for that target was impacted (Figure 4C) (*p* = 0.883, two-sample Kolmogorov-Smirnov test). This indicates that initially, neural activity increased along the neural engagement axes even when doing so negatively impacted task performance. As a result, the initial increase in neural engagement made T2 targets more difficult than they would have been otherwise (relative to the average neural activity produced during Block 1), while T1 targets were made easier.

### Gradual changes in neural engagement led to distinct types of learning

We saw that changes in neural engagement on the first trials using the new BCI mapping occurred regardless of the impact on performance. We wondered whether, given repeated practice with the new mapping over subsequent trials, changes in neural engagement might interact with learning-driven changes for each type of target.

We visualized how cursor velocities under the second mapping changed throughout learning, as a function of whether the initial increase in neural engagement increased (T1) or decreased (T2) the speed of the cursor towards the target (Figure 5A-B). For both types of targets, neural activity on the first trial jumped out abruptly along the neural engagement axis (Figure 5A-B, white circles have moved along the orange arrows relative to the gray circles). Then, over tens of trials, velocities gradually aligned with the target direction, leading to increased speeds towards the target (Figure 5A-B, projection of the blue and red traces increases along the target direction). Were these behaviorally beneficial changes to velocity driven by target-specific changes in neural engagement? We measured the levels of neural engagement for each target during Block 2 after accounting for any changes due to learning by neural reassociation (Golub et al., 2018) (see Methods). In agreement with what we observed earlier (Figure 2D, third panel), we found that neural engagement gradually decreased throughout Block 2 (Figure 5C). Importantly, this decrease in neural engagement was likely beneficial to T2 targets, the ones initially impaired by the increase in neural engagement. In fact, neural engagement decreased more for T2 targets (Figure 5C, red trace) than for T1 targets (Figure 5C, blue trace). This suggests that, as learning proceeded, changes along the neural engagement axis were driven by two components, one target-invariant (neural engagement decreased throughout learning for both target types), and one target-specific (neural engagement decreased by different amounts depending on the target type). As we will show next, these differential changes to neural engagement during learning impacted how quickly performance improved for the two types of targets.

**Figure 5.**
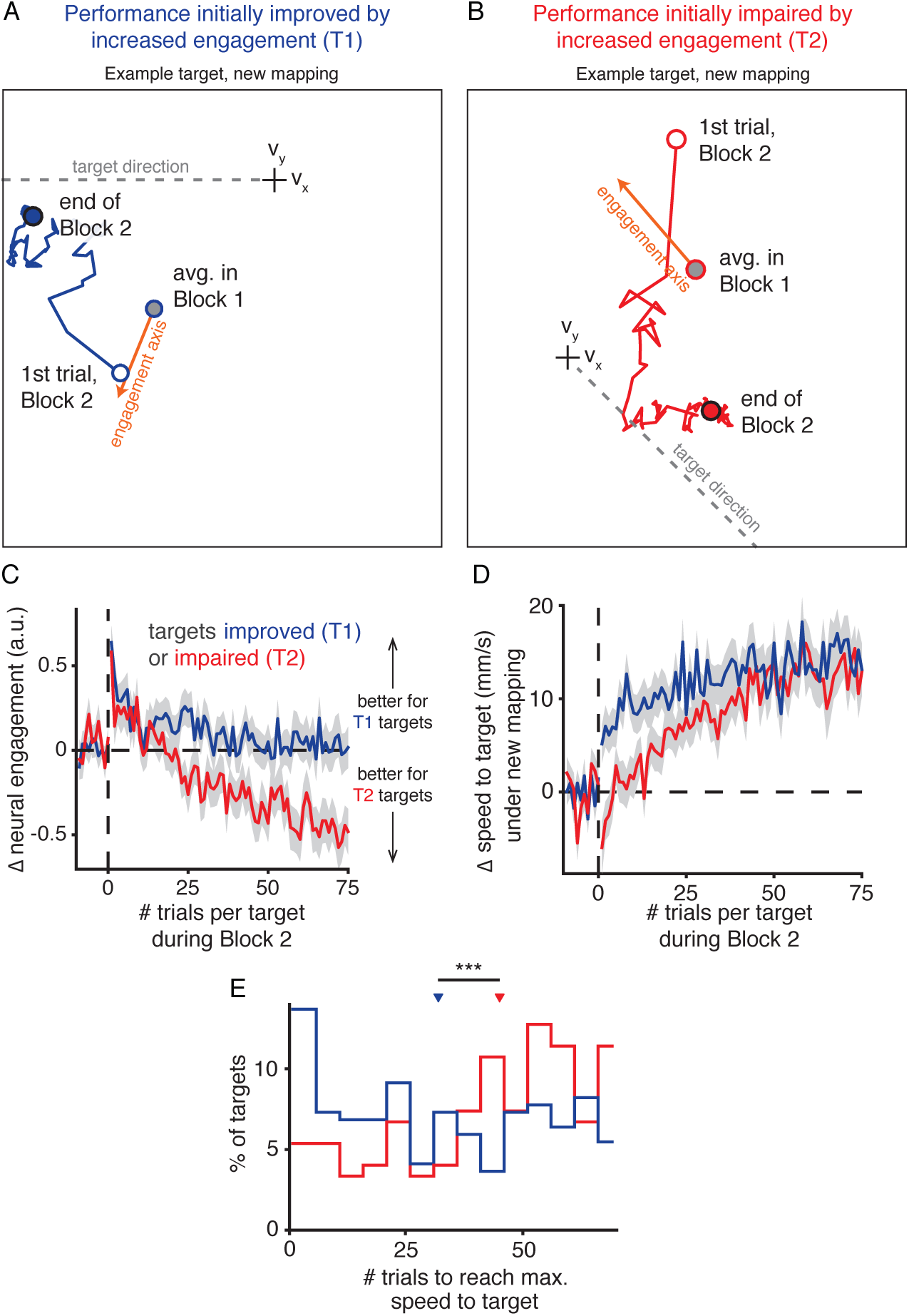
Neural engagement helped to explain why some targets were learned more quickly than others. **A.** Average cursor velocities under the new mapping across trials during Block 2, for an example target (180°, J20120528) where an increase in neural engagement initially improved performance relative to the average activity produced during Block 1 (gray circle). Same conventions as Figure 4A. The blue line depicts how the average velocity evolved throughout Block 2, starting with the first trial to that target (white circle) and ending with the average during the last trials (blue circle). Velocities gradually moved towards the target direction, both decreasing angular error and increasing the speed in the target direction, indicating learning. **B.** Same as A, but for a different example target (315°, J20120601) where an increase in neural engagement was initially expected to impair performance under the new mapping. **C.** Changes along the neural engagement axis during Block 2, averaged across targets (mean ± SE), where targets were split by whether increased neural engagement was expected to initially improve (‘T1’, blue) or impair (‘T2’, red) performance under the new mapping. Trial index is relative to the start of Block 2 for each target. **D.** Changes in cursor speed towards the target under the new mapping during Block 2, relative to the expected speed under the new mapping based on the average neural activity produced during Block 1. Same conventions as C. **E.** Distribution of the number of trials at which each target attained its maximum performance (see Methods), for all T1 and T2 targets. Medians of the two distributions (blue and red triangles) were significantly different (*p* < 0.001, two-sided Wilcoxon rank-sum test).

To quantify the amount of learning for each target, we measured cursor speeds towards the target relative to the speeds monkeys would experience if they continued to use the neural activity they produced prior to the introduction of the new BCI mapping (Figure 5D; see Methods). On the first trial of Block 2, the cursor speed towards the target increased for T1 targets (Figure 5D, blue trace, trial 1), and decreased for T2 targets (Figure 5D, red trace, trial 1). This is in agreement with monkeys immediately increasing neural engagement at the start of Block 2, regardless of its impact on performance (Figure 4C). As Block 2 continued, performance for both target types gradually improved (Figure 5D, blue and red traces, trials 1-75), indicating learning.

Interestingly, monkeys attained their best performance levels more quickly for T1 targets than for T2 targets (Figure 5E; *p* < 0.001, two-sided Wilcoxon rank-sum test). This was not due to a difference in learning rate, as the learning rates for the two target types were not statistically different (*p* = 0.202, two-sided Wilcoxon rank-sum test; see Methods). Additionally, performance levels at the end of Block 2 for the two target types were not statistically different (*p* = 0.957, two-sided Wilcoxon rank-sum test). These results suggest that, although both types of targets were eventually able to achieve similar levels of performance, the initial increase in activity along the neural engagement axes gave performance for T1 targets a “head start,” allowing monkeys to attain their best performance more quickly for T1 targets than for T2 targets. This explanation is at apparent odds with the fact that neural engagement decreased throughout learning for both targets (Figure 5C), which should have led to slower cursor speeds for the T1 targets. In the next section we explore how the initial performance improvements for T1 targets were maintained even as neural engagement decreased throughout learning.

### Neural engagement changed differently in neural dimensions aligned with the new BCI mapping

We have seen how throughout learning, performance for both types of targets gradually improved, regardless of the impact of neural engagement on cursor speeds (Figure 5D). This is in apparent contradiction with the fact that neural engagement decreased throughout learning, even for the targets where decreased neural engagement should have resulted in decreased cursor speeds (i.e., compare the blue traces in Figure 5C and Figure 5D). Crucially, our measurement of neural engagement does not account for which changes in neural engagement affect cursor movements, and which changes do not affect cursor movements. We therefore decomposed each neural engagement axis into two components (Figure 6A; see Methods), where the first component was output-null to the new BCI mapping (i.e., changes in this direction would not impact cursor velocities under the new mapping), and the other component was output-potent (Kaufman et al., 2014; Stavisky et al., 2017b; Hennig et al., 2018). This resulted in measures of output-null and output-potent neural engagement, which allowed us to look specifically at whether neural engagement changed differently depending on whether or not it impacted cursor movements. Changes along the output-null component of the neural engagement axis had no impact on cursor velocities (Figure 6B), and followed the same pattern as the total neural engagement (Figure 5C). By contrast, changes along the output-potent component of the neural engagement axis moved in the directions necessary to yield performance improvements for each target type (Figure 6C). In particular, neural population activity for T1 targets remained elevated along the output-potent component of the neural engagement axis, where performance was initially improved by the increase in neural engagement (Figure 6C, blue trace). This indicates that the net decrease in total neural engagement throughout learning was not entirely agnostic to task performance, as neural activity remained elevated specifically in the neural dimensions that were relevant to controlling the cursor.

**Figure 6.**
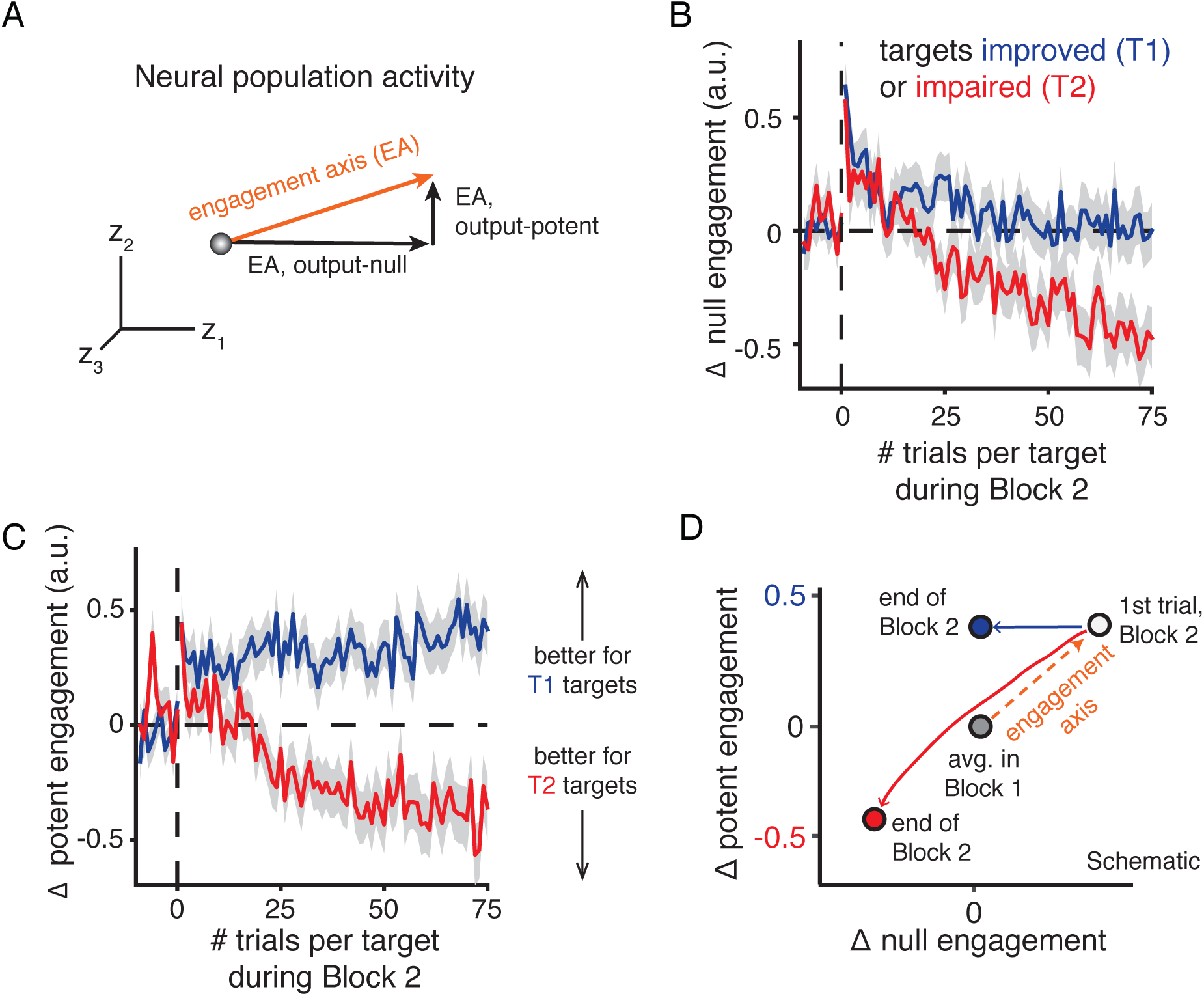
Neural engagement changed differently in output-potent versus output-null dimensions of the new BCI mapping. **A.** Schematic of decomposing a neural engagement axis (EA, orange arrow) into output-null and output-potent components. Given the new BCI mapping, this axis can be decomposed into output-null and output-potent axes, such that only changes in neural activity along the output-potent axis will affect cursor velocities under the new mapping. **B**-**C.** Changes in neural activity along the output-null (B) and output-potent (C) neural engagement axes during learning. Changes in output-null neural engagement do not affect cursor movements, while changes in output-potent neural engagement do. Same conventions as Figure 5C. **D.** Schematic summarizing how neural activity changed during learning for both target types. Circles depict average activity levels before the introduction of the new mapping (gray), on the first trial of Block 2 (white circle), and at the end of learning (red and blue circles) depending on whether neural engagement was predicted to initially improve (blue) or impair (red) performance under the new mapping.

Taken together, these results explain how learning proceeded differently depending on the impact of neural engagement on cursor movements (Figure 6D), resulting in monkeys reaching their best performance levels more quickly for some targets than for others. On the first trial of Block 2, neural activity increased along the neural engagement axis, regardless of its impact on performance (Figure 6D, white circle). This led to immediate performance improvements for T1 targets and decrements for T2 targets (Figure 5D, trial 1). As the trials continued, neural activity gradually decreased along the neural engagement axis for both types of targets (Figure 6D, blue and red arrows). For T2 targets, this decrease in neural engagement was beneficial to performance, yielding progressively faster cursor speeds towards the target. For these targets, neural activity decreased similarly along the components of the neural engagement axis that were output-potent and output-null to cursor velocities under the new BCI mapping (Figure 6D, red arrow). By contrast, for T1 targets, neural activity decreased along the output-null components of the neural engagement axis, but maintained the initial increase in the output-potent components (Figure 6D, blue arrow). This allowed the immediate performance improvements from the increase in neural engagement on trial 1 to be maintained, even as total neural engagement decreased. This resulted in monkeys improving their performance more quickly for T1 targets than for T2 targets.

These results indicate that during learning, neural population activity did not change gradually from the activity observed before learning (Figure 6, ‘avg. in Block 1’) to the activity at the end of learning (Figure 6, ‘end of Block 2’). Rather, neural population activity underwent an abrupt change at the start of learning, improving performance for some targets and impairing performance for others. While the performance levels at the end of learning were similar for both types of targets (Figure 5D), the manner in which neural population activity changed to achieve this performance was quite different (Figure 6D). These findings help to explain why some targets were learned more quickly than others.

## Discussion

We have shown that large, trial-to-trial fluctuations in M1 population activity along neural engagement axes exhibited hallmarks of an arousal- or motivation-like process. While monkeys learned a new BCI mapping, neural activity increased abruptly along neural engagement axes on the first trial of learning, regardless of its effect on behavioral performance. This indicates that changes in neural activity during learning need not be a gradual transition between the activity produced prior to learning and the activity produced at the end of learning. On subsequent trials during learning, neural activity retreated along neural engagement axes, which interacted with learning. This led monkeys to learn some targets more quickly than others, based on how neural engagement axes related to behavior. Thus, changes in internal states can interact with the learning process and influence how quickly different task goals are learned.

In this study, we found that trial-to-trial changes in neural engagement were positively correlated with changes in the monkey’s pupil size, a common psychophysical index for an animal’s internal state (Beatty, 1982; McGinley et al., 2015; Joshi et al., 2016). The term ‘internal state’ is used broadly, but typically refers to any neural signal that does not directly reflect, but may interact with, sensory encoding or behavior generation (McGinley et al., 2015). This includes internal states related to computation (e.g., internal models (Shadmehr and Holcomb, 1997), reward prediction (Schultz et al., 1997), working memory (Courtney et al., 1997)), but also those reflective of more autonomic processes (e.g., arousal (Vinck et al., 2015), motivation (Mazzoni et al., 2007), task engagement (Steinmetz et al., 2019)). We have termed the internal state identified in the present work ‘neural engagement’ because its stereotyped time course was suggestive of changes in the monkey’s engagement with the task throughout the experiment (e.g., increases in neural engagement following pauses in the experiment and the introduction of a new BCI mapping). This is reminiscent of, but potentially distinct from, the concept of ‘task engagement’ (Otazu et al., 2009; Steinmetz et al., 2019), referring in this case to the difference between an animal actively versus passively experiencing a stimulus. While our current study design does not allow us to identify the exact source of changes in neural engagement, in the following paragraphs we consider multiple possibilities and how they might explain (or fail to explain) the results in the present work.

*Is it intended speed?* Neural engagement may be related to, but is likely distinct from, the monkey’s intended movement speed. Neurons in M1 have long been known to reflect movement speed (Georgopoulos et al., 1986; Schwartz and Moran, 1999). We observed that during arm movements, increased neural engagement predicted increased hand speed towards the target (Figure S3). This raises the possibility that neural engagement may simply reflect the monkey’s intended movement speed. However, during BCI learning, we observed a gradual decrease in neural engagement during repeated trials to the same target, despite the fact that performance for many targets would have been improved by maintaining this increased neural engagement (Figure 5C). Therefore, if neural engagement simply reflected intended movement speed, it would be necessary to explain why monkeys would intend to move slower when doing so would reduce their reward rate. One possible explanation might be that the monkey’s intended movement speed is modulated by an internal state such as motivation or reward expectation. In fact, studies of “movement vigor,” measured behaviorally as the reaction time and/or peak movement speed during eye or reaching movements (Mazzoni et al., 2007; Xu-Wilson et al., 2009; Dudman and Krakauer, 2016; Yttri and Dudman, 2018; Sedaghat-Nejad et al., 2019; Shadmehr et al., 2019), have found that the vigor (or speed) with which we execute a movement is not constant over time, but varies depending on context. Movement vigor is therefore thought to reflect a cost-benefit analysis, such that vigor increases when there is a higher subjective utility (e.g., expected reward) for doing so (Shadmehr et al., 2019). Consistent with this prediction, neural engagement was higher at the start of the experiment, and following pauses in the experiment (Figure 2D); in both cases, the resumption of the experiment indicates to the monkey a higher expectation of reward, because completing trials resulted in a reward. However, we also saw an increase in neural engagement following the introduction of the new BCI mapping, a time when the monkey’s reward expectation should be lower, given that the new BCI mapping will immediately decrease his reward rate. Thus, increases in neural engagement do not always reflect increased reward expectation, suggesting that neural engagement may not simply reflect movement vigor.

*Is it a feedback response?* Previous work has established that M1 population activity reflects sensory feedback following a perturbation, for both mechanical (Pruszynski et al., 2011, 2014; Omrani et al., 2014, 2016) and purely visual (Stavisky et al., 2017b) perturbations. At first glance, these results may appear similar to our observation of an immediate increase in neural engagement following the introduction of a new BCI mapping. However, our results differ in two key ways. First, while we did find a fast increase in neural engagement (within a single trial), neural engagement then decreased gradually over subsequent trials (Figure 2D). It is not known from these previous studies whether the magnitude of the sensory feedback signal should decay over subsequent trials (nor would we expect this to be the case). Second, neural engagement followed a similar time course during the portion of each trial before cursor feedback was available (Figure 2D, last subpanel), indicating that this signal was not directly reflecting visual feedback. Thus, neural engagement does not simply reflect sensory feedback. More recent work has indicated the presence of another fast, within-trial response to an unexpected mechanical perturbation (Crevecoeur et al., 2019). In this study, humans performed reaches with a manipulandum, where an unexpected force was applied to subjects’ arms on randomly selected trials. The experimenters observed that, after a perturbed trial, reaches on subsequent unperturbed trials showed increased hand speeds towards the target, and co-contraction of the arm muscles, consistent with a theory of robust control (Başar and Bernhard, 2008). While it is possible that such a theory may explain the quick increase in neural engagement following events that were unpredictable to the subject (Figure 2D), this theory also predicts that movement speeds and co-contraction should *increase* on subsequent trials using the new BCI mapping (if the new mapping is akin to a force perturbation) (Crevecoeur et al., 2019), in contrast to the subsequent decrease in neural engagement that we observed in the present study.

*Is it arousal?* Recent work identified a slowly varying correlate of internal state in the neural population activity of prefrontal cortex and visual area V4 while monkeys performed a perceptual decision-making task (Cowley et al., 2020). The authors present evidence that this “slow drift” in population activity reflected an arousal or impulsivity signal, which biased animals’ decisions. The authors propose that this signal may arise from the release of a neuromodulator such as norepinephrine (NE), distributed by the locus coeruleus (LC) (Aston-Jones and Cohen, 2005; McGinley et al., 2015). We speculate that the neural engagement signal identified in the present work may have a similar origin. This would also be consistent with recent work in rodents reporting brain-wide modulation associated with behavioral variables such as facial expression (Stringer et al., 2019) and licking (Stringer et al., 2019; Allen et al., 2019) that can indicate changes in arousal. What might be the role of an arousal signal, if any, in M1? It has been proposed that the LC signals uncertainty in the environment (Yu and Dayan, 2003; Sales et al., 2019), and that the release of NE modulates a trade-off between explorative-exploitative behaviors (Aston-Jones and Cohen, 2005). From this perspective, the increases in neural engagement that we observe following pauses in the experiment and at the start of learning may be due to the phasic release of NE by the LC. If these changes indeed serve a function, such as indicating a change in the environment or driving exploration, our results suggest that this response is relatively coarse or stereotyped across task goals, because the increase in neural engagement persisted even when it caused detriments to behavior.

Our results add to a growing list of work finding population-level signatures of internal state fluctuations (Cohen and Maunsell, 2010; Ecker et al., 2014; Rabinowitz et al., 2015; Lin et al., 2015; Williamson et al., 2016; Huang et al., 2019; Stringer et al., 2019; Allen et al., 2019; Cowley et al., 2020). While these changes need not adversely impact stimulus encoding (Averbeck et al., 2006; Moreno-Bote et al., 2014) or downstream readout (Hennig et al., 2018; Perich et al., 2018; Semedo et al., 2019), empirically these changes can be correlated with measurable deficits in behavior (Ruff and Cohen, 2019; Cowley et al., 2020). In our work, knowing the causal relationship between neural population activity and behavior (i.e., via the BCI) allowed us to directly assess how changes in neural activity impacted behavioral performance. We leveraged this knowledge to establish that M1 population activity underwent large-variance changes during learning even when these changes were detrimental to behavioral performance. Thus, internal state fluctuations can impact not only concurrent behavior, but also future behavior due to their interaction with learning.

## Methods

### Experimental details

Experimental methods are described in detail in Sadtler et al. (2014) and Golub et al. (2018). Briefly, we recorded from the proximal arm region of primary motor cortex (M1) in three male rhesus macaques using implanted 96 electrode arrays (Blackrock Microsystems). All animal care and handling procedures conformed to the NIH Guidelines for the Care And Use of Laboratory Animals and were approved by the University of Pittsburgh’s Institutional Animal Care and Use Committee. Data from monkeys J and L were first presented in Sadtler et al. (2014), while data from monkey N were first presented in Golub et al. (2018). We recorded from 85 to 94 neural units in each session. The activity of each neural unit is defined as the number of threshold crossings recorded by an electrode in non-overlapping 45 ms bins. The average firing rate of the neural units across sessions was 46 ± 7, 38 ± 8, and 56 ± 13 spikes/s (mean ± s.d.) for monkeys J, L, and N, respectively.

During each experimental session, a monkey performed an eight-target center-out task by modulating his recorded neural activity to control the velocity of a computer cursor on a screen. Each session involved two different BCI mappings. The first ‘intuitive’ mapping was chosen to provide the monkey with proficient control of the cursor. The animal used the intuitive mapping for 321 ± 96 trials (mean ± s.d.), after which the mapping was switched abruptly to a second, new BCI mapping that the monkey had never controlled before. This new mapping was chosen so as to be initially difficult for the monkey to use, and the monkey was given 698 ± 227 trials (mean ± s.d.) to learn the new mapping. Both BCI mappings were chosen so that they were controlled exclusively by the neural activity within the monkey’s intrinsic manifold (defined below).

At the beginning of each trial, a cursor appeared in the center of the workspace, followed by the appearance of one of eight possible peripheral targets (chosen pseudo-randomly among *θ* ∈ { 0°, 45°, 90°, 135°, 180°, 225°, 270°, 315°}). For the first 300 ms of the trial, the velocity of the cursor was fixed at zero. After this, the velocity of the cursor was controlled by the animal through the BCI mapping. If the animal acquired the peripheral target with the cursor within 7.5 s, he received a water reward, and the next trial began 200 ms after target acquisition. Otherwise, the trial ended, and the animal was given a 1.5 s time-out before the start of the next trial.

During each experiment we monitored the monkey’s pupil diameter (arbitrary units) using an infrared eye tracking system (EyeLink 1000; SR Research, Ottawa, Ontario). The eye tracker was first turned on while monkeys used the intuitive mapping, but this time varied from session to session. Pupil diameter was always measured while monkeys controlled the new BCI mapping.

#### Selecting the BCI mappings

Each session began with the monkey performing a block of calibration trials, as described in Sadtler et al. (2014). The calibration procedure for monkey J involved either passive observation of cursor movement, or closed-loop BCI cursor control using the previous day’s BCI mapping. For monkeys L and N, we used a closed-loop calibration procedure that gradually stepped from passive observation to closed-loop control. We first z-scored the spike counts recorded during these calibration trials, where z-scoring was performed separately for each neural unit. We then applied factor analysis (FA) to the z-scored spike counts to identify the 10D linear subspace (i.e., the ‘intrinsic manifold’) that captured dominant patterns of co-modulation across neural units (Santhanam et al., 2009; Churchland et al., 2010; Harvey et al., 2012; Williamson et al., 2016; Athalye et al., 2017; Huang et al., 2019). The factor activity, **z**_*t*_ ∈ ℝ^10×1^, was then estimated as the posterior expectation given the z-scored spike counts, **u**_*t*_ ∈ ℝ^*q*×1^, where *q* is the number of neural units: 

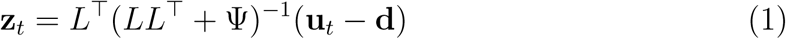

Above, *L*, Ψ and **d** are FA parameters estimated using the expectation-maximization algorithm, where *L* is termed the *loading matrix*, and Ψ is constrained to be a diagonal matrix. The factor activity, **z**_*t*_, can be interpreted as a weighted combination of the activity of different neural units. We refer to **z**_*t*_ as a “population activity pattern.”

As discussed above, each experiment consisted of animals using two different BCI mappings. Each BCI mapping translated the resulting moment-by-moment factor activity (**z**_*t*_) into a 2D cursor velocity (**v**_*t*_) using a Kalman filter: 

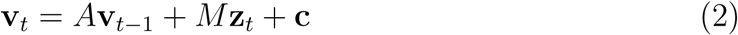

For the ‘intuitive’ BCI mapping, *A* ∈ ℝ^2×2^, *M* = *M*_1_ ∈ ℝ^2×10^, and **c** ∈ ℝ^2×1^ were computed from the Kalman filter parameters, estimated using the calibration trials. For the second, ‘new’ BCI mapping, we changed the relationship between population activity and cursor movement by randomly permuting the elements of **z**_*t*_ before applying Equation 2. This permutation procedure can be formulated so that Equation 2 still applies to the second BCI mapping, but for a new matrix *M*_2_ ∈ ℝ^2×10^ used in place of *M*_1_ (Sadtler et al., 2014).

We orthonormalized **z**_*t*_ so that it had units of spike counts per time bin (Yu et al., 2009). This was done by finding an orthonormal basis for the columns of the matrix *L* above. We can do this by applying the singular value decomposition, yielding *L* = *USV*^*T*^, where *U* ∈ ℝ^*q*×10^ and *V* ∈ ℝ^10×10^ have orthonormal columns and *S* ∈ ℝ^10×10^ is diagonal. Then, we can write 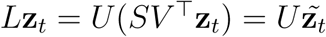. Because *U* has orthonormal columns, 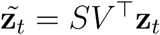 has the same units (spike counts per time bin) as **u**_*t*_. For notational simplicity, we refer to 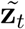 as **z**_*t*_ throughout.

Note that the data analyzed in this study were part of a larger study involving learning two different types of BCI mapping changes: within-manifold perturbations (WMP), described above, and outside-manifold perturbations (OMP) (Sadtler et al., 2014). We found that animals learned WMPs better than OMPs, and so we only analyzed WMP sessions in this study. In total, we analyzed data from 46 WMP sessions; this consisted of 25 sessions from monkey J, 10 sessions from monkey L, and 11 sessions from monkey N.

#### Hand control experiments

Data were collected from a fourth monkey for three sessions. During these experiments, the monkey performed an eight-target center-out task by moving his hand to control a computer cursor. An infrared marker was taped to the back of the monkey’s hand and tracked optically using an Optotrak 3020 system. The marker position was used to update the position of the cursor in real-time on a stereoscopic computer monitor. During these experiments we recorded from the proximal arm region of primary motor cortex (M1) using an implanted 96 electrode array (Blackrock Microsystems).

Similar to the BCI control experiments, the targets shown on each trial were chosen pseudo-randomly. At the beginning of each trial, a target (sphere; radius: 6 mm) was presented in the center of the reaching workspace. The animal was trained to move the cursor (sphere; radius: 6 mm) to this start target and hold for 0-100 ms. A peripheral target (sphere; radius: 6 mm) was presented at the end of this hold period. Water reward was delivered if the target was acquired within 1.5 s and the cursor was held on the target for a random hold period drawn uniformly from 150-550 ms. The next trial was initiated 200 ms after the trial ended, regardless of success or failure. The data analyzed includes 160 trials of baseline center-out trials, where the marker position was directly mapped to the cursor position, followed by 320 trials where a visuomotor rotation was applied to all reaches (40° CW, 40° CCW, and 30° CW for the three sessions, respectively).

To match the analysis procedure used in the BCI experiments, we took spike counts in non-overlapping 50 ms bins, and z-scored the spike counts using the mean and standard deviation of each neural unit during baseline reaches. We then applied factor analysis to the z-scored spike counts recorded during all baseline reaches to identify a 12D linear subspace, where 12 was the number of dimensions that maximized the cross-validated log likelihood. We then orthonormalized the resulting 12D factor activity. All analyses of population activity considered only these top 12 factors.

### Data analysis

#### Time step selection

In the BCI experiments, spike counts were taken in non-overlapping 45 ms bins (‘time steps’), indexed here by *j* = 1, *…, T*, where *T* is the number of time steps in a given trial, and *j* = 1 is the time step where the target first appeared. Each trial consisted of three intervals of interest: 1) the pre-target interval (*j* ≤ 2, or 90 ms), during which the monkey had not yet perceived the target due to sensory processing delays; 2) the freeze interval (*j* ≤ 6), during which the cursor was frozen in place at the center of the workspace; and 3) the cursor control interval (*j* ≥ 7), where the cursor velocity was determined by Equation 2. Unless otherwise noted, all analyses used data only during the cursor control interval.

We noted that when the cursor was near the target, or at the end of long trials, cursor movements were often idiosyncratic (e.g., reflecting small corrective movements), and so we discarded from our analyses any time steps where the cursor was more than 65% of the way to the target, and any time steps *j* > 20. To report trial-averaged quantities, we wanted to ensure that all neural activity within the same trial came from time steps where the monkey attempted to push the cursor in the same direction. This was especially important given that we compared the time course of neural engagement during learning on a target-by-target basis (see Figure 4, Figure 5, and Figure 6). We therefore analyzed only the time steps where the angle between the cursor and target was within 22.5° of the target direction on that trial. Performing our analyses without this exclusion criterion did not change our results.

We analyzed both correct and incorrect trials in this study. We reasoned that sufficiently large increases in neural engagement (e.g., on the first trial using the new BCI mapping) may slow down the cursor’s speed to the extent that the monkey is unable to obtain the target. Removing incorrect trials would then bias any analyses that compare levels of neural engagement between targets whose performance was improved versus impaired by neural engagement (see Figure 4, Figure 5, and Figure 6).

For the hand control experiments, we analyzed data from the 15 time steps of each trial immediately following the appearance of the target (which cued the monkey to begin moving his hand towards the target).

#### Quantifying behavior

To relate changes in neural engagement to the monkey’s ability to improve his performance using the new BCI mapping (Figure 5), we assessed both the monkey’s performance and neural engagement on a moment-by-moment basis (i.e., for each time step within a trial). To quantify the monkey’s moment-by-moment behavior, we calculated the speed to the target contributed by a given neural activity pattern under the new BCI mapping (i.e., “cursor progress” defined in Golub et al. (2018)). Specifically, given the neural activity pattern **z**_*j*_ produced at a particular time step *j*, the new BCI mapping parameters *M*_2_ and **c** (see Equation 2), and a unit vector **p**_*j*_ pointing from the cursor position at time step *j* to the target position, we computed the speed to the target, *s*_*j*_ (shown in Figure 5D), as: 

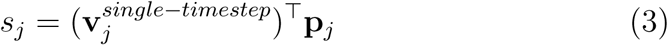

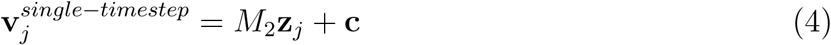

where 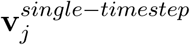 is the velocity contributed by the neural population activity **z**_*j*_ recorded at a single time step (Golub et al., 2018) (i.e., ignoring the contribution from the neural population activity at previous time steps; see Equation 2). Assessing performance in this manner ensures that our measures of neural engagement and performance (as in Figure 5) are both assessed using precisely the same neural activity.

Let *s*_*θ*_(*t*) be the speed to the target under the new BCI mapping for a given target *θ*, on trial *t* ∈ {1, *…, T*_*θ*_} during Block 2 (i.e., *s*_*θ*_(*t*) is the average of *s*_*j*_ for all time steps *j* from trial *t*). To report average performance changes during Block 2 (Figure 5D), we averaged *s*_*θ*_(*t*) for each *t* across all T1 targets and T2 targets separately. To find the trial at which performance for each target *θ* was maximized (Figure 5E), we first found the running mean of *s*_*θ*_(*t*) in a sliding eight trial window. Let 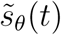 be the resulting running mean (defined for 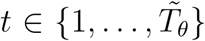, where 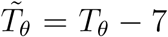 due to the smoothing). The trial at which performance for each target *θ* was maximized was then arg max_*t*_ 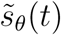. To test whether performance levels at the end of Block 2 differed between T1 and T2 targets, we used 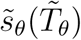 as the performance level of target *θ* at the end of Block 2. Finally, to assess whether learning rates differed between T1 and T2 targets, for each *θ* we fit a saturating exponential to *s*_*θ*_(*t*) with free parameter *τ* > 0: 

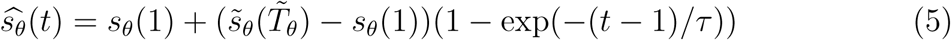

where *τ* is the learning rate, governing how quickly *s*_*θ*_(*t*) transitions from initial performance, *s*_*θ*_(1) (unsmoothed because *s* changed more quickly early in learning), to performance at the end of Block 2, 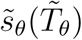. For each target, *τ* was chosen so as to minimize the mean squared error between 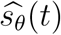 and *s*_*θ*_(*t*) for all *t*.

#### Identifying neural engagement axes

For each experimental session (for either BCI or hand control), we sought to identify a set of *neural engagement* axes, one per target direction, capturing the dimensions along which neural activity varied in the absence of learning pressure (i.e., while monkeys used the intuitive BCI mapping, or during baseline reaches, respectively). For each target *θ*, we defined the neural engagement axis, **a**_*θ*_ ∈ ℝ^10^, with ∥**a**_*θ*_ = 1∥, as the direction of greatest variance in the factor activity recorded during all trials to that target. Identifying this direction in the factor activity rather than in the spiking activity ensures that we focus on the shared covariance among neural units rather than variance that is independent to each unit.

The neural engagement axes are sign-invariant, which would ordinarily prevent us from identifying ‘positive’ versus ‘negative’ changes along these vectors. This would make averaging values of neural engagement across sessions and targets meaningless, because ‘positive’ values of neural engagement for one session or target might not correspond to ‘positive’ values of neural engagement on a different session or target. However, we observed that the neural engagement axes involved the activity of nearly all neural units changing in the same direction (Figure S2). This allowed us to choose the sign of **a**_*θ*_ in a consistent manner, by ensuring that positive values of neural engagement corresponded to increases in the firing rate for the majority of units. This allowed us to average across values of neural engagement across targets and sessions, as presented in the main text.

#### Quantifying neural engagement

As described above, we identified the neural engagement axes, **a**_*θ*_, for each target *θ* ∈ {0°, 45°, 90°, 135°, 180°, 225°, 270°, 315°} during Block 1, while monkeys controlled the intuitive BCI mapping. We also identified the mean neural activity, 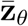, produced for each target. We noted that, although monkeys showed proficient control of the intuitive BCI mapping, there were still substantial fluctuations around 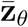. We estimated these fluctuations along **a**_*θ*_, which we term *neural engagement*. For the neural activity, **z**_*j*_, observed at a given time step *j* for target *θ*, we estimated neural engagement, or *e*_*j*_, as follows:

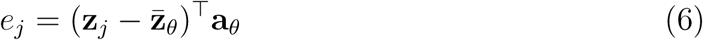

We estimated neural engagement for each time step of the experiment, and then averaged the values across all time steps within a given trial (as shown in Figure 2, Figure 4, and Figure S1). To combine these values across trials to different targets (as plotted in Figure 2 and Figure S1), we then z-scored the neural engagement for each target separately, using the mean and standard deviation of the neural engagement measured during the last 10 trials to each target during Block 1.

##### Inferring changes in neural engagement during learning

To estimate neural engagement during Block 2, we cannot simply use Equation 6, because some of the changes in neural activity across trials will also be due to learning (e.g., by neural reassociation (Golub et al., 2018)). According to neural reassociation (Golub et al., 2018), to move the cursor in a particular direction *θ* ∈ [0, 2*π*) during Block 2, the monkey samples the neural population activity he used for movements in a potentially *different* direction *θ*′ ∈ [0, 2*π*) during Block 1. Thus, to estimate neural engagement during Block 2 (as shown in Figure 5 and Figure 6), we used the following: 

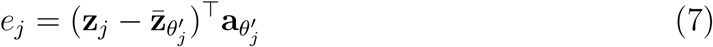

where 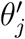 is no longer necessarily equal to the target direction, *θ*. We estimated 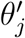 from the neural activity, **z**_*j*_, which is reasonable provided that changes in neural activity due to *θ*_*j*_ and *e*_*j*_ are not entirely overlapping. We therefore estimated 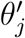 by finding the direction that the cursor would have moved if **z**_*j*_ were produced under the intuitive mapping, as changes in neural engagement tended to have less effect on the cursor’s movement direction using the intuitive mapping. This procedure allowed our estimate of *θ*′ to vary as the monkey learned to control the new BCI mapping, thus factoring out any changes in neural activity due to neural reassociation. To compute 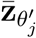 and 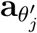 for any continuous value of 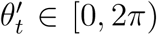, we used a cubic spline to interpolate between the values measured for each *θ* ∈ {0°, 45°, 90°, 135°, 180°, 225°, 270°, 315°}.

For this procedure we again estimated neural engagement at each time step, and then averaged the values across all time steps within a given trial. As described above, we z-scored the neural engagement for each target separately. The z-scoring again used the mean and standard deviation of the neural engagement measured during the last 10 trials to each target during Block 1, ensuring that values of neural engagement could be compared across different blocks of the experiment.

In the above procedure, the neural engagement axes corresponding to a given *θ* are assumed to be the same during both Block 1 and Block 2. We confirmed that the neural engagement axes estimated before learning (during Block 1) and after learning (at the end of Block 2) were similar (Figure S5), indicating that the largest fluctuations in neural activity occurred along similar dimensions throughout the experiment.

##### Comparing neural engagement to pupil size

For each session, we estimated the correlation between the estimated neural engagement with the monkey’s pupil size (Figure 2E-F). Pupil sizes were measured consistently only during Block 2 (see *Experimental details* above), and so this analysis used trials only from Block 2, for all sessions where Block 2 consisted of at least 200 trials (45 of 46 sessions). To compare slow timescale fluctuations between neural engagement and pupil size during Block 2, we first applied boxcar smoothing to the trial-averaged measurements of each quantity with a sliding window of 30 trials. We then computed the Pearson’s correlation between the two time series.

##### Variance explained by changes in neural engagement

We sought to estimate the amount of variance in the neural population activity due to changes in neural engagement (Figure 2G). To estimate the variance for trials to a given target, we first found the neural activity **z**_*t*_ for each trial to that target, along with the corresponding neural engagement, *e*_*t*_. The measure of the variance explained by changes in engagement for that target was then 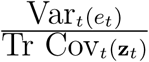. To compute the *total* amount of variance explained by changes in engagement, we computed the same metric above, but used the activity from all trials combined rather than just the trials to a particular target.

##### Predicting the impact of neural engagement on performance under the new mapping

We estimated the impact of increased neural engagement on cursor movements under the new mapping for each target. To do this, we quantified the predicted change in the cursor speed to the target given an increase in neural activity along the positive direction of the neural engagement axis. Specifically, let 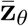 be the average neural activity recorded during Block 1 for target *θ*, and let **a**_*θ*_ be the corresponding neural engagement axis. Then we labeled that target as improved by an increase in neural engagement if we expected the speed to target to increase (see Equation 3):

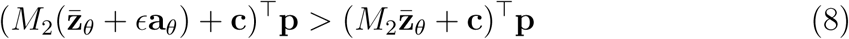

where *M*_2_ and **c** are the parameters of the new BCI mapping, and *ϵ* > 0. This procedure was used to identify the targets for which performance would initially be improved versus impaired by an increase in neural engagement, as introduced in Figure 4.

##### Identifying output-potent and output-null engagement axes

Given a neural engagement axis, **a** ∈ ℝ^10^, not all changes in neural activity along this axis will lead to changes in cursor velocity through the new BCI mapping, *M*_2_. This is because the mapping between neural activity and cursor velocity, given by Equation 2, is a linear mapping from 10D to 2D, implying that *M*_2_ has a non-trivial null space, *Nul*(*M*_2_). To identify which components of **a** will result in changes in cursor velocity, we can find bases for the null space, *Nul*(*M*_2_), and the row (or potent) space, *Row*(*M*_2_) (Hennig et al., 2018). To do so, we took a singular value decomposition of *M*_2_ = *USV*^*T*^, with *U* ∈ ℝ^2×2^, *S* ∈ ℝ^2×10^, and *V* ∈ ℝ^10×10^, where the columns of *S* were ordered so that only the first two columns had non-zero elements. Then, we let *R* ∈ ℝ^10×2^ be the first two columns of *V*, and *N* ∈ ℝ^10×8^ be the remaining eight columns. The columns of *N* and *R* are mutually orthonormal and together form an orthonormal basis for the 10-dimensional space of factor activity. This allows us to rewrite the neural engagement axis for each target *θ* as the sum of a null-engagement axis, 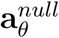, and a potent-engagement axis, 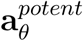: 

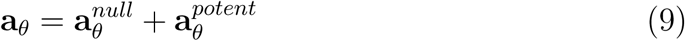

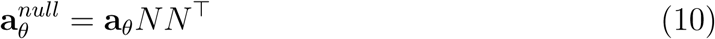

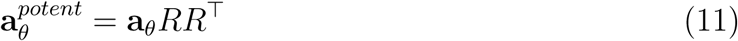

We then normalized 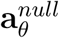 and 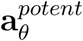 to be unit vectors. The resulting axes were used to compute values of null and potent engagement, as shown in Figure 6, by using these axes in Equation 7.

## Acknowledgments

The authors would like to thank Jessica Graves, Benjamin Cowley, Matthew Smith, and Eric Yttri for helpful discussions. This work was supported by the Richard King Mellon Presidential Fellowship (JAH), Carnegie Prize Fellowship in Mind and Brain Sciences (JAH), NIH R01 HD071686 (APB, BMY, and SMC), NSF NCS BCS1533672 (SMC, BMY, and APB), NSF CAREER award IOS1553252 (SMC), NIH CRCNS R01 NS105318 (BMY and APB), NSF NCS BCS1734916 (BMY), NIH CRCNS R01 MH118929 (BMY), NIH R01 EB026953 (BMY), Simons Foundation 543065 (BMY), and Pennsylvania Department of Health Research Formula Grant SAP 4100077048 under the Commonwealth Universal Research Enhancement program (SMC and BMY).

## Author Contributions

J.A.H. performed the analyses. M.D.G., P.T.S., K.M.Q., A.P.B., S.M.C., and B.M.Y. designed the animal experiments. E.R.O., L.A.B., and P.T.S. performed the animal experiments. E.R.O., S.I.R., and E.C.T.-K. performed the animal surgeries. J.A.H., A.P.B., S.M.C., and B.M.Y. wrote the manuscript. All authors discussed the results and commented on the manuscript. A.P.B., S.M.C., and B.M.Y. contributed equally to this work.

---

**Figure S1.**
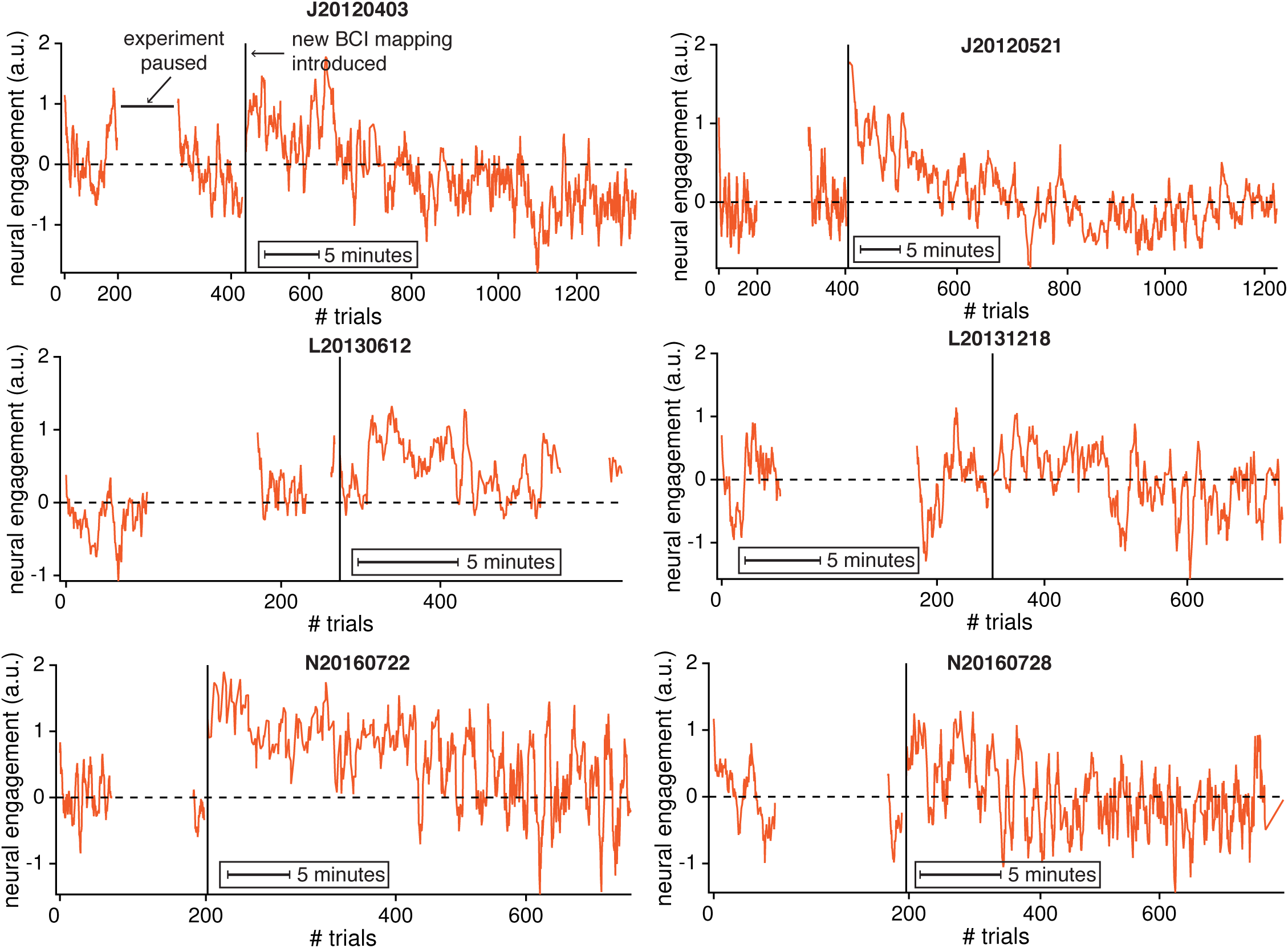
Neural engagement showed stereotyped changes relative to experimental events in multiple example sessions from three monkeys. Same conventions as Figure 2C.

**Figure S2.**
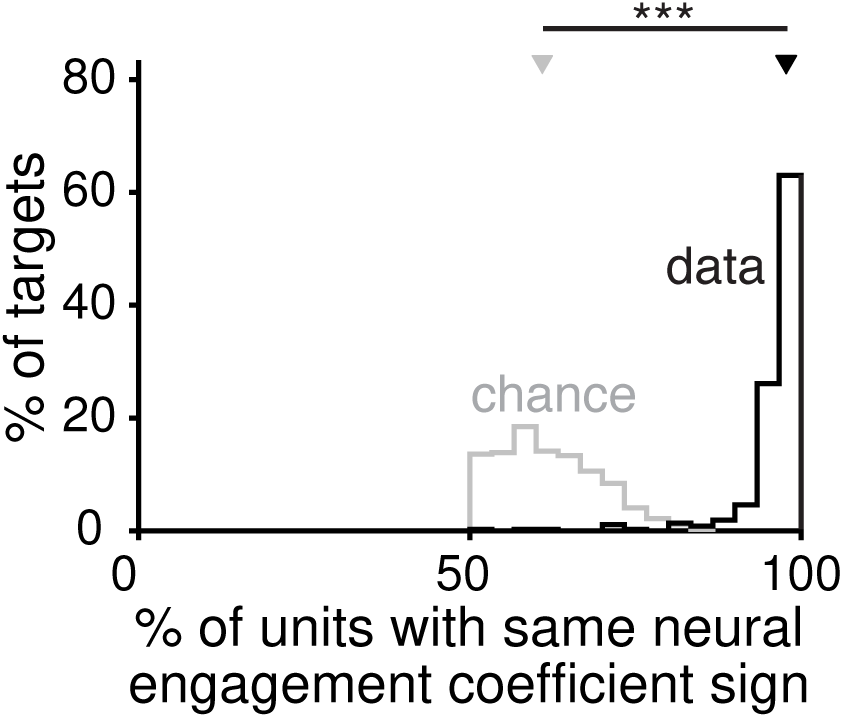
Changes in neural engagement corresponded to nearly all neural units increasing or decreasing their activity together. We wanted to understand how changes in neural engagement were represented by the activity of individual units. For each target, a neural engagement axis was defined in 10-dimensional factor space. We used the *q* × 10 loading matrix from factor analysis (see Methods) to define the neural engagement axis in the *q*-dimensional population activity space of the *q* recorded units. For example, if there were 90 units, the neural engagement axis would have 90 coefficients, describing how changes in neural engagement for a given target would be represented by the activity of each of the 90 units. For each target, we computed the percentage of units whose coefficients had the same sign (for whichever sign was in the majority, so that percentages could never be below 50%). Shown in black is the distribution of these percentages across the neural engagement axes for all targets across all sessions (bootstrapped 95% C.I. [97.6%, 97.7%]). For reference, in gray, is the distribution after sampling random dimensions in factor space, and computing the corresponding effects on individual neural units (bootstrapped 95% C.I. [59.7%, 62.5%]). Triangles depict the medians of the ‘data’ and ‘chance’ distributions, which were significantly different (*p* < 0.001, two-sided Wilcoxon rank-sum test).

**Figure S3.**
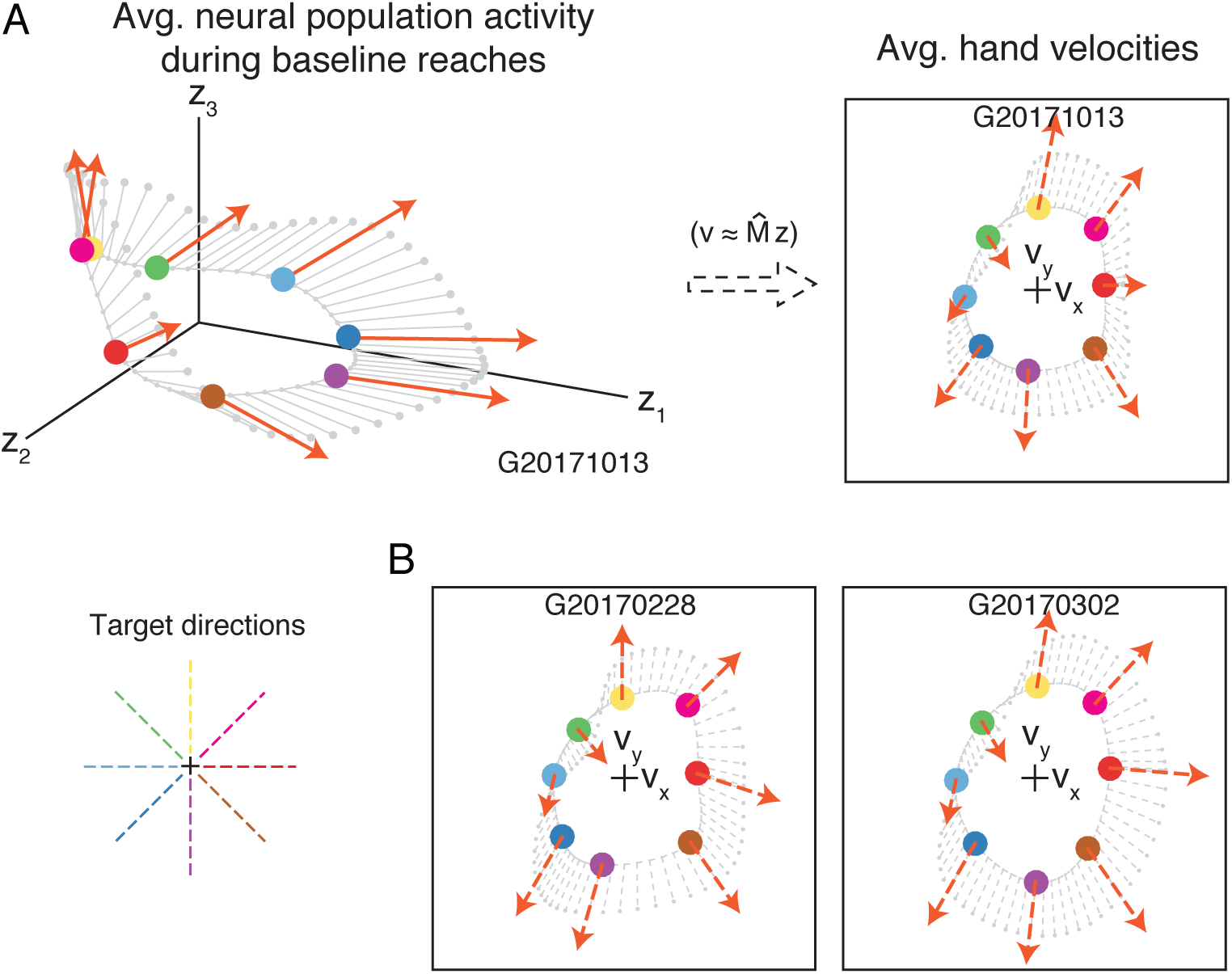
Increased neural engagement during arm movements predicted faster hand speeds towards most targets. **A.** For the experiments involving arm movements (see Methods), we visualized the average neural population activity (circles, left subpanel) and neural engagement axes (orange arrows, left subpanel) during baseline reaches to each of eight targets. Same conventions as Figure 3B. We also visualized the monkey’s average hand velocity during reaches to each target (circles, right subpanel). Unlike during BCI control, we do not know the causal relationship between neural population activity and hand velocity. To understand how changes in neural engagement related to hand velocity, we used linear regression to predict the monkey’s hand velocity during baseline reaches at each 50 ms time step during the movement epoch of every trial, using the neural population activity recorded 100 ms prior. Cross-validated *r*_2_ for the x- and y- components of hand velocity were 67% and 77%, respectively. The linear regression model 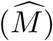 allowed us to estimate how increases in the neural engagement related to the monkey’s average hand velocity towards each target (orange dashed arrows), and to intermediate target directions (gray dashed arrows). In this session, an increase in neural engagement predicted an increase in the monkey’s hand speed towards all but the 135° target. This suggests that differences in the neural engagement axes across targets may have behavioral relevance. ‘Target directions’ panel is a legend depicting the color corresponding to each target direction. **B.** We repeated the above procedure during the other two arm movement sessions. Across sessions, increases in neural engagement predicted faster hand speeds towards all but the 135° target.

**Figure S4.**
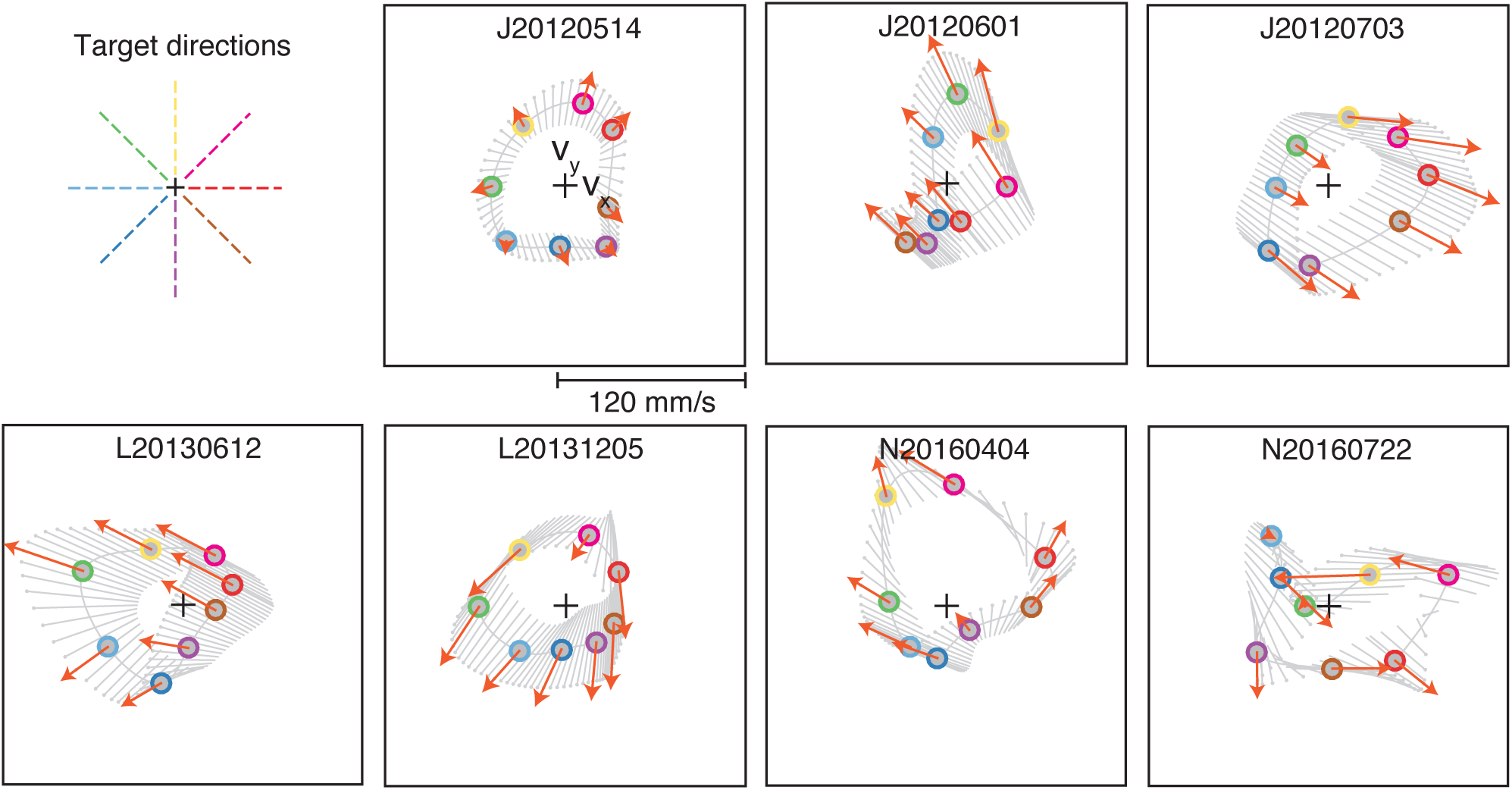
New BCI mappings induced a variety of relationships between neural engagement and cursor velocity, across targets and sessions. Same conventions as the bottom right panel of Figure 3B, for multiple example sessions (all with the same scale). ‘Target directions’ panel is a legend depicting the color corresponding to each target direction.

**Figure S5.**
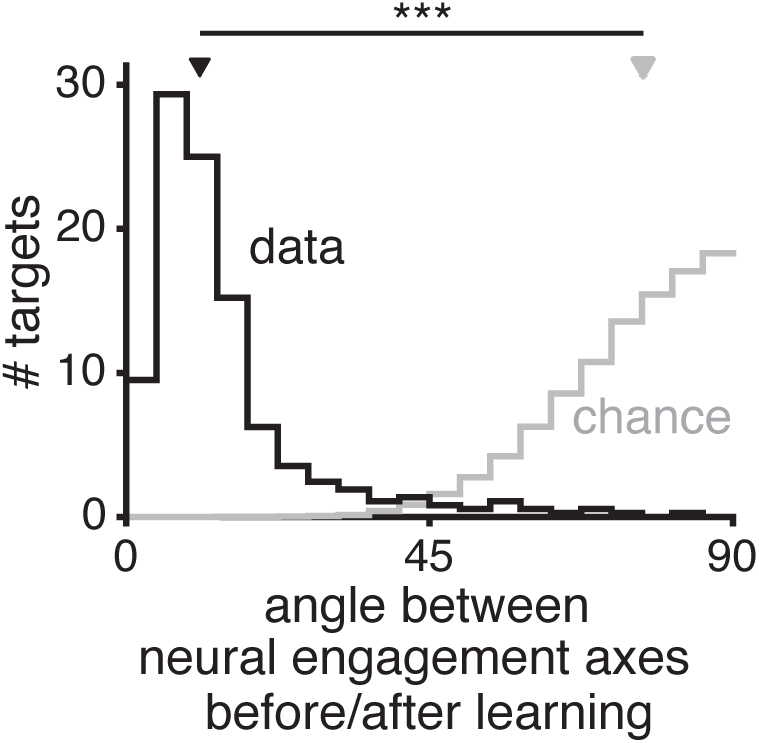
Neural engagement axes were largely unchanged after learning. Distribution of the angle (‘data’, in black) between the neural engagement axis identified for each target during Block 1 (‘before learning’) vs. during the last 50 trials of Block 2 (‘after learning’). To identify neural engagement axes during the last 50 trials of Block 2, we used the same procedure as used during Block 1 (i.e., the procedure used in the main text; see Methods), but applied to the last 50 trials of Block 2. ‘Chance’ (in gray) indicates the distribution of the angle between random directions in ten-dimensional space. Triangles depict the medians of the ‘data’ and ‘chance’ distributions, which were significantly different (*p* < 0.001, two-sided Wilcoxon rank-sum test).

